# Single-chain lanthanide luminescence biosensors for cell-based imaging and screening of protein-protein interactions

**DOI:** 10.1101/2020.03.10.985739

**Authors:** Ting Chen, Ha T. Pham, Ali Mohamadi, Lawrence W. Miller

**Affiliations:** Department of Chemistry, University of Illinois at Chicago, Chicago IL, USA

## Abstract

Research tools that enable imaging or analysis of protein-protein interactions (PPIs) directly within living cells provide unique and valuable biological insights and can also aid drug discovery efforts. Here, we present lanthanide-based, Förster resonance energy transfer (lanthanide-based FRET, or LRET) biosensors for time-gated luminescence (TGL) imaging or multiwell plate analysis of PPIs. Polypeptide chains comprised of an alpha helical linker flanked by a Tb(III) complex, GFP and two binding domains exhibit large differences in long-lifetime, Tb(III)-to-GFP LRET-sensitized emission between open (unbound) and closed (bound) states. We used TGL microscopy to image ca. 500% increases in Tb(III)-to-GFP LRET following rapamycin addition to NIH 3T3 cells that expressed biosensors bearing FKBP12 and the rapamycin binding domain of m-Tor (FRB) at each terminus. Much larger signal changes, up to ca. 2500%, were observed when cells were grown in 96-well or 384-well plates and analyzed using a TGL plate reader. We also measured the interaction of p53 and HDM2 and its inhibition within intact HeLa cells grown in 96-well plates and estimated a z’-factor of 0.5 for the assay. The modular design and high dynamic range of Tb(III)-based LRET biosensors will facilitate versatile imaging and cell-based screening of PPIs.

## INTRODUCTION

Fluorescence-based methods that enable the imaging or analysis of protein-protein interactions (PPIs) directly in living cells are critical tools for fundamental biological research and for drug discovery.^*1*^ PPIs regulate nearly all biological processes, and cell-based methods of study are required because PPIs are often weak, occur transiently in pairs or larger complexes, exist within large and overlapping biochemical networks or exert their function only when sequestered in distinct sub-cellular locations.^*2*^ Mechanistic studies benefit from live-cell microscopy with genetically encoded fluorescent proteins (FPs) that can capture the spatial and temporal dynamics of PPIs relative to cells’ response to stimuli or changes in phenotype.^*3, 4*^ Efforts to discover drugs that inhibit or activate PPIs are aided by cell-based screens and counter-assays that utilize multi-well plate readers and that can be used to evaluate hits or leads for cytotoxicity, membrane permeability or off-target effects.^*5-7*^

PPIs are most commonly imaged in cells using FP-based biosensors that rely on the phenomenon of Förster resonance energy transfer (FRET) to transduce biochemical events into changes in fluorescence intensity, wavelength or lifetime.^*1, 4*^ FRET is non-radiative, dipole-dipole energy transfer from a donor fluorophore to a nearby (usually closer than 10 nm) acceptor species that has an absorption spectrum which overlaps the donor’s emission spectrum.^*3*^ FRET-based imaging of PPIs in live cells can be achieved with a so-called dual-chain biosensor configuration by expressing the binding partners as genetic fusions to appropriately paired FP donors and acceptors such as cyan and yellow (CFP, YFP) or green and red (GFP, RFP). Interaction of the fusion proteins results in an increase in FRET which may be observed as a reduction in the emission intensity or lifetime of the donor and a concomitant increase in donor-sensitized, acceptor emission. Alternatively, single-chain biosensor designs are constructed such that target analyte binding or the interaction of two affinity domains induces a conformational change that modulates intramolecular FRET efficiency between donor and acceptor.^*4*^

While FRET can be a powerful tool for single-cell, microscopic imaging, FRET biosensor signal changes are often subtle. Consequently, FRET-based assays are less commonly used for medium-throughput or high-throughput screening (HTS) applications where large signal changes and low variability are desirable. Nevertheless, some FRET-based screening methods as well as those based on bioluminescence resonance energy transfer (BRET) have been reported.^*8-11*^ Non-FRET, cell-based assays for PPI discovery that have been adapted to a high-throughput rate of analysis include methods based on sub-cellular redistribution of fluorescently labeled proteins (suitable for high-content imagers),^*12, 13*^ reporter fragment complementation assays (e.g., split GFP, split luciferase),^*14*^ and reporter gene hybrid-like systems,^*15, 16*^ and some methods based on FRET or bioluminescence resonance energy transfer (BRET).^*8, 9, 11*^ However, all of these available cell-based PPI assays suffer from one or more limitations, including low signal-to-background ratio (SBR) or dynamic range, high rates of false positives/negatives, or protein sequestration at non-physiologic sites.

Here, we present lanthanide-based FRET (LRET) biosensors for live cell imaging and multiwell plate analysis of PPIs. These sensors incorporate luminescent Tb(III) complexes with ms-scale excited state lifetimes as FRET donors and GFP as acceptors and are amenable to time-gated luminescence (TGL) detection. With TGL, pulsed, near-UV light is used to excite the specimen, and long-lived Tb(III) or Tb(III)-to-GFP LRET signals are captured after a brief delay (∼µs) occurs during which ns-scale sample autofluorescence and directly excited acceptor fluorescence decays (**Figure 1A**). We characterized sensor performance using two model systems: i) the rapamycin-induced interaction between FK binding protein 12 (FKBP12) and the rapamycin binding domain of m-Tor (FRB);^*17*^ and ii) the therapeutically relevant interaction between p53 and HDM2.^*18*^ Our single-chain biosensor design incorporated a rigid alpha-helical linker sequence comprised of multiple repeats of four glutamic acid or arginine residues alternated with four lysine residues (ER/K) flanked by EGFP and *Escherichia coli* dihydrofolate reductase (eDHFR). The affinity binding elements were positioned at the N- and C-termini of the sensors (**Figure 1B**). The eDHFR domain binds with high specificity and affinity (*K*_*D*_, ∼1 nM) to heterodimers of trimethoprim linked to a luminescent Tb(III) complex,^*19*^ permitting selective labeling of the sensor construct.

**Figure 1.**
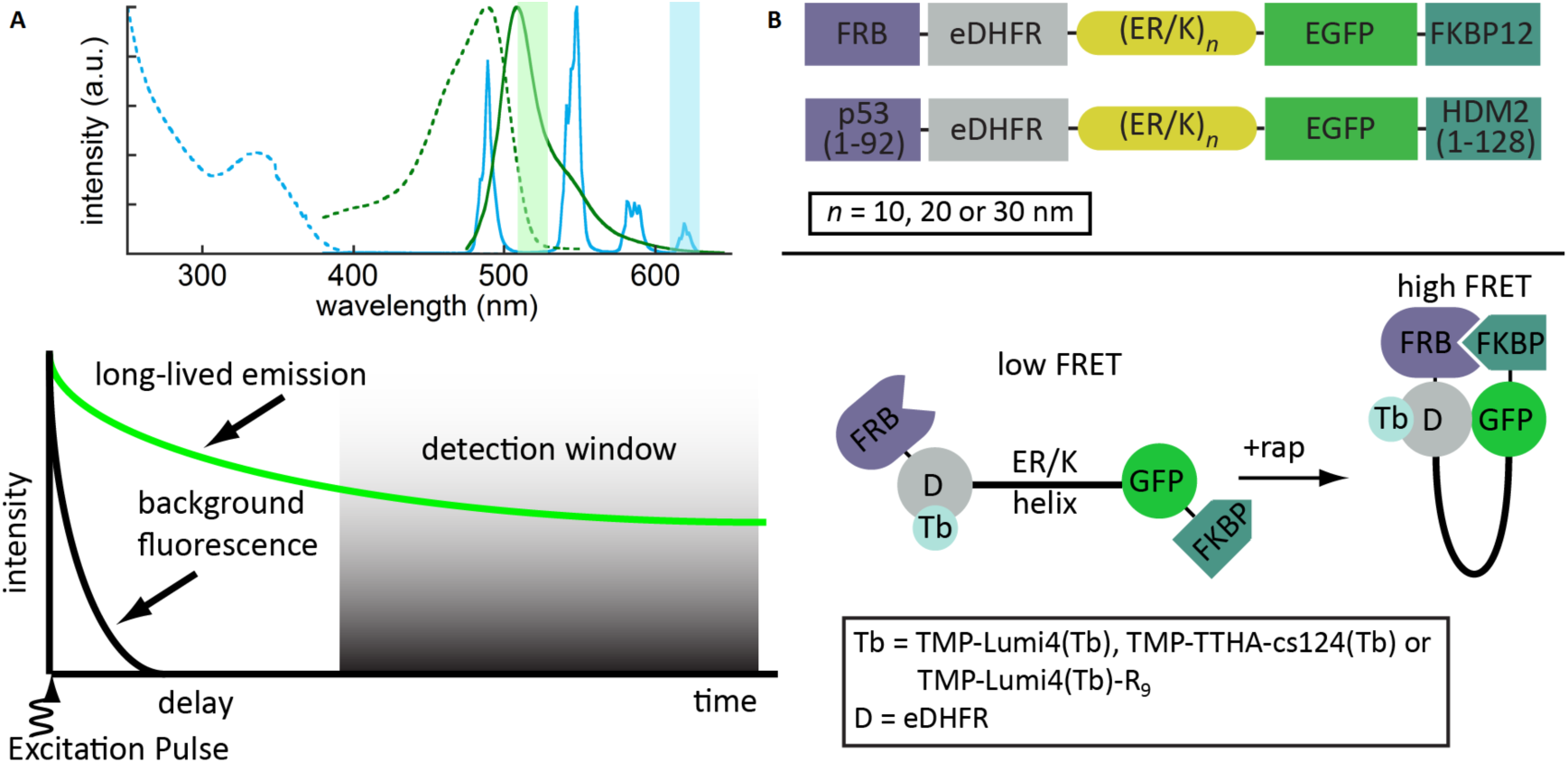
Single chain, Tb(III)-based FRET biosensor design leverages the narrow, multi-line emission spectrum and ms-scale excited state lifetime of Tb(III) complexes to facilitate high signal-to-background, time-gated luminescence (TGL) detection. **A)** (Top) Excitation (dotted) and emission (solid) spectra of Tb(III) (cyan) and GFP (green). Colored bars show emission bandpass for detecting Tb(III) and Tb(III)-to-GFP LRET signals. (Bottom) Insertion of a delay (10 µs) between pulsed excitation and detection enables background-free detection of Tb(III) luminescence and Tb(III)-to-GFP LRET-sensitized emission. **B)** Biosensor design (Top). A ER/K helix motif (length 10, 20 or 30 nm) separates LRET partners and affinity pairs. (Bottom) Absent interaction, the ER/K helix maintains affinity and detection elements far apart, ensuring low baseline LRET signal. Stochastic breaking of helix linker permits close approach and binding of affinity domains.

Overexpression of the biosensors in HeLa or NIH3T3 fibroblast cells followed by TGL microscopy or TGL analysis in 96-well and 384-well plates enabled sensitive imaging and detection of biosensor activity. Remarkable sensor dynamic ranges of over 500% and over 2500% were observed for rapamycin-induced activation of FKBP12/FRB interaction in live cell microscopic images and in 96-well plates, respectively. Statistically robust detection of FKBP12/FRB interaction or inhibition and p53/HDM2 inhibition was observed in 384-well plates. The high performance seen here with model systems and a modular sensor design indicate that Tb(III)-based, single-chain FRET biosensors can be applied to analyze a wide variety of PPIs.

## RESULTS AND DISCUSSION

Several noise sources reduce the dynamic range and precision of FRET measurements and hinder the ability to monitor two or more FRET pairs in a single specimen. These noise sources include overlapped donor and acceptor excitation and emission spectra, non-specific background from sample autofluorescence or directly excited acceptor emission and (for imaging applications) differences in the local concentrations of donor-labeled and acceptor-labeled proteins. Consequently, intensity-based FRET microscopy requires acquisition of two or three images at different wavelengths to separate biochemically relevant FRET signals from non-FRET fluorescence background and to normalize the FRET signal to the amounts of donor or acceptor. When imaging single-chain biosensors, two-color, ratiometric measurements of the FRET signal and either donor or acceptor signals is often sufficient to quantify FRET changes.^*3*^ High levels of non-specific background (e.g., from library compound fluorescence) have limited the development of live-cell, FRET-based HTS using FPs or other conventional fluorophores,^*9*^ although the recent development of fluorescent lifetime plate readers has increased measurement precision and allows for more robust assay development.^*20*^ The combination of TGL detection and luminescent lanthanide probes with ms-scale excited state lifetimes is used extensively to overcome many of the limitations of conventional FRET for HTS applications,^*21*^ and TGL microscopy has been explored for live cell imaging studies.^*22, 23*^

A key biosensor performance parameter is dynamic range, defined here as the maximum observed difference between the mean, donor-denominated or acceptor-denominated FRET ratios in the On-state and Off-state of the sensor.^*24, 25*^ While dual-chain biosensors typically exhibit a high dynamic range, quantitative analysis of intermolecular FRET is complicated by substantial differences in the local concentrations of donor-labeled and acceptor-labeled proteins. Single-chain biosensors overcome this problem by maintaining a 1:1 donor:acceptor ratio. However, single-chain sensors may fold into an Off-state conformation where the donor and acceptor labels are in close proximity, leading to high baseline FRET and low dynamic range. Dynamic range is also reduced when the sensor adopts an On-state conformation where the orientation of the donor’s and acceptor’s relative dipole moments disfavors FRET. For these reasons, many single-chain biosensors yield FRET ratio changes lower than 50%.^*24, 25*^ Considerable efforts have been made to improve single-chain FRET biosensor dynamic range by using circularly permutated FPs to optimize fluorophore orientation,^*26*^ mutating FPs to increase their inherent dimerization^*11*^ and engineering linker sequences that better separate affinity elements and fluorophores in the low-FRET state.^*25, 27*^

### Biosensor Design

We aimed to develop single-chain, Tb(III)-based biosensors but were concerned about high baseline signals. Dipole orientation doesn’t affect LRET because Tb(III) or Eu(III) emit with multiple dipole moments at many different orientations.^*28*^ However, LRET between lanthanides and organic fluorophores or fluorescent proteins can typically be observed over relatively long distances (10 – 20 nm) compared to conventional FRET.^*29*^ Moreover, during the long excited state lifetime of a lanthanide chromophore, many conformations can be sampled, some of which might bring the donor and acceptor close to one another. In order to minimize baseline LRET signals, we incorporated into our sensor design a semi-rigid α-helix linker sequence, which consists of an alternating sequence of approximately four glutamic acid residues followed by approximately four arginine or lysine residues [(E/R)_4_/K_4_, or ER/K linker). As reported by Sivaramakrishnan and Spudich, the ER/K linker adopts an alpha-helical geometry in solution.^*30*^ However, it was speculated that the ER/K helix can break stochastically, permiting close approach of elements positioned on either end. Thus, insertion of the ER/K sequence between affinity binding elements and FRET partners can yield a biosensor with low baseline FRET because the donor and acceptor are held far apart in the Off-state, yet the ends can still bind one another. Because the affinity elements are tethered together, their effective concentration depends only on linker length and is independent of solution concentration. The overall fraction of an ER/K biosensor in the closed or On-state depends only on the *K*_*D*_ of the affinity elements and linker length. Consequently, protein-protein interactions may be observed and analyzed even when the overall sensor concentration is far below the *K*_*D*_^*31*^.

### TGL microscopy of biosensors in live cells

NIH3T3 fibroblasts were stably transfected with plasmid DNA encoding a single fusion protein under control of a Tet-responsive promoter that contained the following elements (from *N- to C-* terminus): FRB, eDHFR, ER/K, GFP and FKBP12 (Figure 1). Three stably transformed cell lines were created that expressed sensors with ER/K linker lengths of 10 nm, 20 nm or 30 nm. Following overnight induction of protein expression with doxycycline, cells were incubated in culture medium containing a cell-permeable, luminescent Tb(III) complex, (TMP-Lumi4-R_9_,^*32, 33*^ **Figure S1**) (12 µM, 15 min, room temperature), washed with PBS, immersed in imaging medium and then imaged immediately. Steady-state images of GFP fluorescence and time-gated images of Tb(III) luminescence and Tb(III)-to-GFP sensitized emission revealed sensor distribution throughout the cytoplasm and Tb(III) probe distribution throughout the cytoplasm and nucleus (**Figure 2A**). In a time-series image sequence of cells expressing the sensor with a 20 nm linker, the donor-denominated LRET (LRET/Tb) ratio increased to over 300% of its initial value about 15 min after rapamycin addition (**Figure 2B**). Very large increases in both LRET/Tb and the acceptor-denominated LRET ratio (LRET/GFP) were observed in rapamycin-stimulated cells expressing biosensors containing 10 nm, 20 nm, and 30 nm (E/R)_4_/K_4_ linkers (**Fig. 2C**). The dynamic ranges of both LRET/Tb and LRET/GFP signals increased with linker length. In cells expressing FKBP12/FRB biosensors with 10, 20, and 30 nm ER/K linkers, the maximum observed changes in mean LRET/Tb were 87%, 288% and 525%, respectively. The maximum, microscopically observed increases in mean LRET/GFP were 61%, 378% and 470% for linker lengths of 10, 20 and 30 nm, respectively.

**Figure 2.**
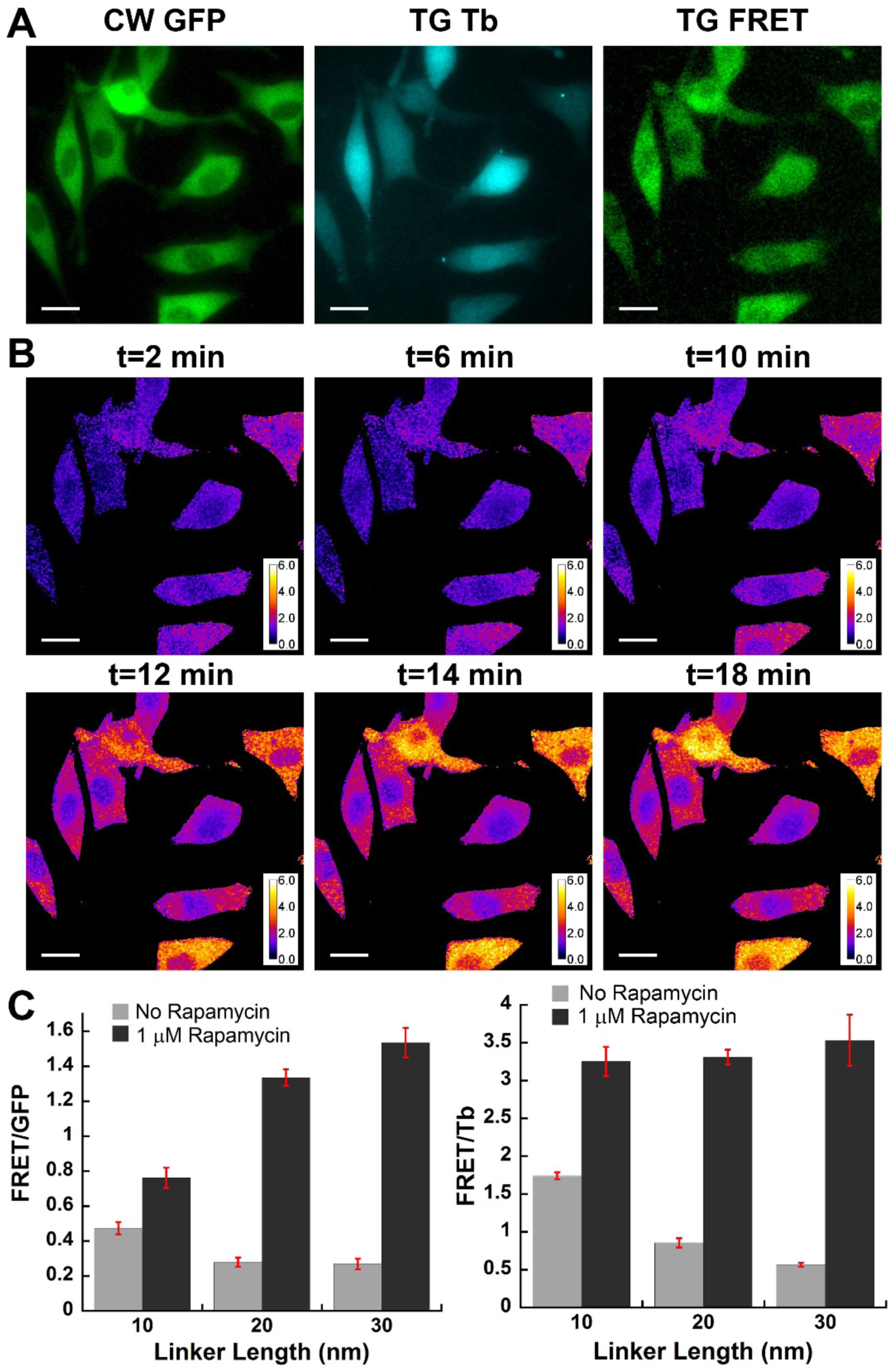
Time-gated luminescence microscopy enables two-channel, ratiometric imaging of LRET PPI biosensors with high dynamic range. **A.** Representative images of NIH3T3 fibroblasts cells stably expressing FRB-eDHFR-(ER/K)_20_-GFP-FKBP12 approximately 20 min after stimulation with rapamycin (1 µM). Micrographs: CW GFP, steady-state GFP fluorescence (λ_ex_, 480 ± 20 nm; λ_em_, 535 nm ± 25 nm); TG Tb, time-gated Tb(III) luminescence (λ_ex_, 365 nm, λ_em_, 620 nm ± 10 nm, gate delay 10 µs); TG LRET, time-gated Tb(III)-to-GFP sensitized emission (λ_ex_, 365 nm; λ_em_, 520 ± 10 nm, gate delay 10 µs). Scale bars, 20 µm. TG Tb and TG LRET channel images were rendered at identical contrast. **B**. Color maps of the same cells shown in **A** depict the ratio of the TG LRET image to the TG-Tb image at various time points following rapamycin stimulation. **C.** Biosensor dynamic ranges increase with the length of ER/K linker due to reduction in baseline, or Off-State LRET signals. Bar graphs depict the mean, pixel-wise LRET/Tb or LRET/GFP ratios measured in regions of interest drawn within cells both before and 25 min after addition of rapamycin. Values given are averaged from 10 or more cells for each condition. Error bars, SEM.

### Detection of PPIs and their Inhibition in Multi-well plates

Often, conventional FRET-based detection of cellular PPIs at medium throughput (96-well plate) or high-throughput (384-well plate) is impossible because of the aforementioned limitations in FRET S/N and dynamic range and the relatively small amounts of protein in each sample well.^*9*^ We sought to assess the potential of our Tb(III) biosensors for detection and quantification of PPIs and their inhibition in multi-well plate format following expression in live mammalian cells. NIH 3T3 cells stably expressing single-chain, FKBP/FRB biosensors (containing 20 nm, or 30 nm ER/K linkers) were seeded into 96-well plates (40,000 cells/well) and grown overnight in medium containing doxycycline to induce protein expression. A rapamycin titration assay was first performed to obtain the optimal rapamycin concentration to induce the FRB/FKBP12 interaction. Lysis buffer containing TMP-Lumi4-Tb^*19*^ (50 nM) and serial dilutions of rapamycin (final conc., 5 µM to 0.5 nM) was added to the wells, and the plate was incubated at room temperature for 15 min.

Following incubation, the time-gated Tb(III)-to-GFP LRET and Tb(III) emission signals were measured at 520 nm and 615 nm, respectively. The 520 nm signal from each well was divided by the 615 nm signal in order to minimize well-to-well variability resulting from differences in probe amounts or sample absorbance. Then, the mean 520/615 emission ratio from 12 control wells containing non-expressing cells and lysis buffer solution (50 nM TMP-Lumi4-Tb, no rapamycin) was subtracted from each sample well to yield a background-corrected, LRET/Tb ratio. A non-linear regression (NLR) fit to a plot of FRET/Tb ratio vs. rapamycin concentration yielded EC_50_ values of 22 ± 2 nM and 18 ± 2 nM for cells expressing biosensors with 20 nm or 30 nm ER/K linkers, respectively. Maximal interaction was observed at rapamycin concentrations equal to or exceeding 500 nM (**Figure S2**).

In order to further assess the performance of our model system, we stimulated sensor-expressing cells grown in either 96-well or 384-well plates with lysis buffer containing 1 µM rapamycin and TMP-TTHA-cs124(Tb)^*19, 34*^ (**Figure S1**) (final conc., 25 nM). Again, the mean LRET/Tb ratio from control wells (n = 16) containing non-expressing cells and lysis buffer with TMP-cs124-TTHA(Tb) but lacking rapamycin was subtracted from the LRET/Tb ratio measured for each sample well. An increase in dynamic range with ER/K linker length was observed in the 96-well plate data, similar to that seen in microscopy data. However, the magnitude of the measured dynamic range was substantially higher. Cells expressing FKBP12/FRB biosensors with 10 nm, 20 nm, or 30 nm ER/K linkers exhibited dynamic ranges of 165%, 1700%, and 2500%, respectively. For all sensor constructs, the maximum observed LRET/Tb ratio was similar. However, the sensor with 10 nm ER/K linker had a higher baseline LRET signal (**Figure 3A**).

**Figure 3.**
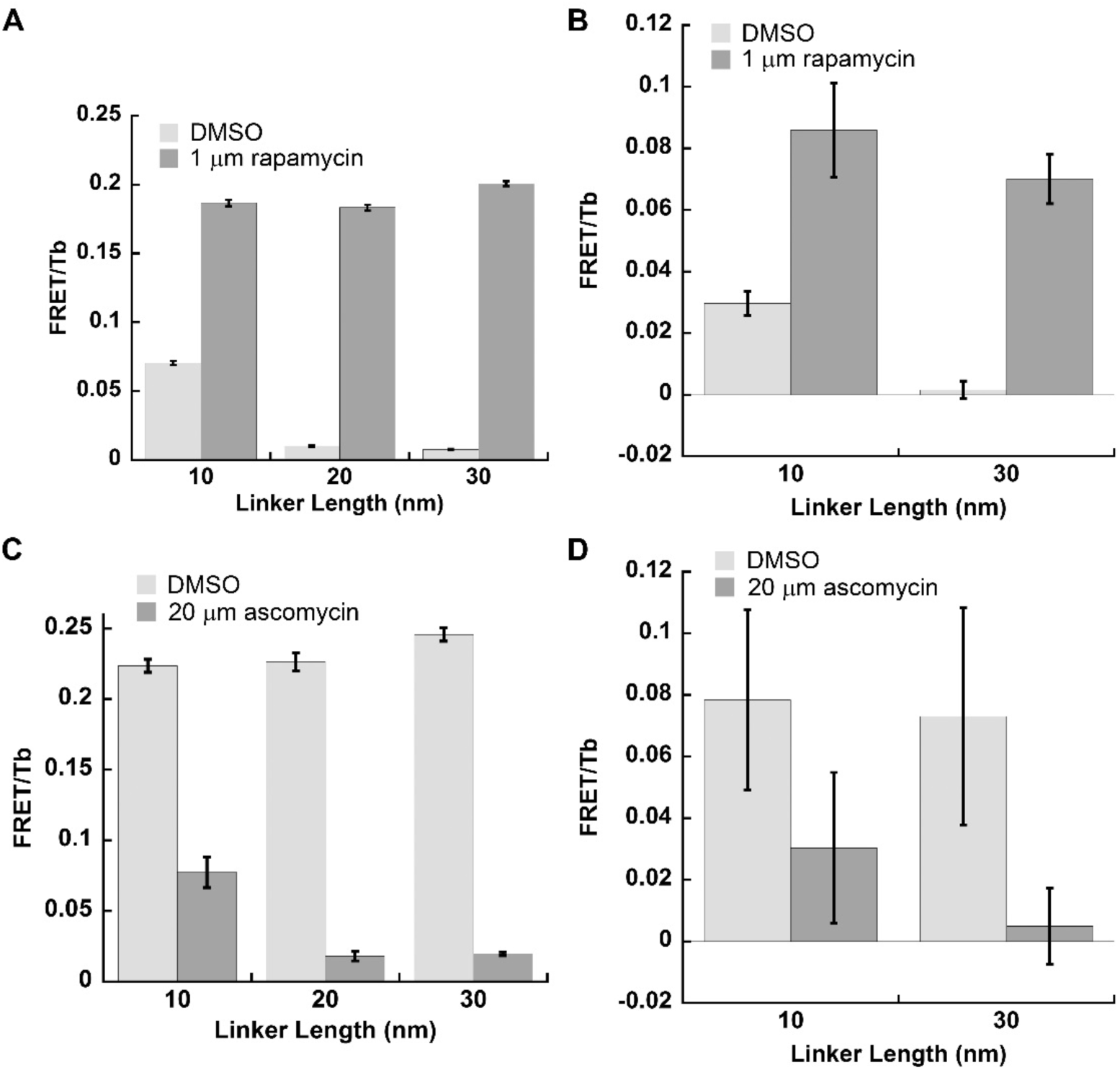
TGL analysis robustly detects FKBP12/FRB interaction and its inhibition following permeabilzation of sensor-expressing cells grown in multiwell plates. NIH 3T3 fibroblast cells expressing FKBP/FRB sensors with varied ER/K linker lengths were seeded into 96-well (**A, C**) or 384-well (**B, D**) plates at cell densities of 40,000 or 8,000 cells/well, respectively. Following overnight incubation, cells were treated with lysis buffer containing TMP-TTHA-cs124 (25 nM). Time-gated emission signals (gate delay, 0.2 ms) at 520 nm and (Tb-to-GFP LRET) and 620 nm (Tb only) were measured using a time-resolved fluorescence plate reader. **A, B.** LRET/Tb ratios were substantially larger when lysis buffer contained rapamycin (1 µM). **C, D.** Cells were treated with lysis buffer containing TMP-TTHA-cs124 (25 nM) and rapamycin (0.33 uM). Time-gated signals were then measured as in (**A**,**B**). Addition of ascomycin (20 µM) to lysis buffer decreased LRET/Tb emission ratios by more than 60% for all sensor linker lengths in both 96-well and 384-well plates. Bar graphs depict mean LRET/Tb ratio measured for positive controls (n = 16) and for negative controls (n = 8). Error bars, SD.

We further evaluated our system by calculating the Z’-factor, a standard quality metric for HTS assays.^*35*^ Z’-factor is calculated from the standard deviations and means of the maximum and minimum observed signal levels obtained with positive and negative controls (i.e., without library compounds present) according to eqn. 1.

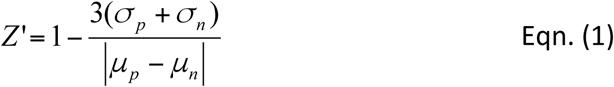

*Z’* can vary between -∞ and 1, with values > 0.5 considered to be a very good assay, values between 0 and 0.5 considered marginal, and < 0 an unacceptable assay.^*35*^ For measurement of FKBP12/FRB biosensor activation following cell permeabilization in 96-well plates, Z’ ranged from 0.72 to 0.89 for all sensors. While these results clearly indicate a highly robust assay, high-throughput assays require the capability to measure at least 100,000 compounds per day, and this requires analysis in 384-well plates. In 384-well plates, we obtained a relatively poor *Z’* factors about zero for the sensor with a 10 nm ER/K linker and a value of 0.41 for the sensor with a 30 nm ER/K linker (**Figure 3B**). Some variance in the data may be attributable to manual plate preparation, suggesting further improvement in assay robustness may be achievable with optimization.

In order to further evaluate the potential of Tb(III) biosensors for multi-well plate applications, we measured the effects of ascomycin as an inhibitor of the rapamycin-induced, FKBP12/FRB interaction^*36*^. Ascomycin was titrated against a constant concentration of rapamycin (0.333 µM) in permeabilized cells expressing the FKBP/FRB sensors with either 20 or 30 nm ER/K linker lengths. Non-linear fitting yielded IC_50_ values of 0.39 ± 0.05 µM for both sensors (**Figure S2**). Full inhibition with 20 µM ascomycin yielded LRET/Tb signal decreases of more than 60% for all ER/K Linker lengths (**Figure 3C-D**). While Z’ factors greater than 0.69 were obtained for all 96-well plate assay conditions, large relative error yielded negative Z’ values for ascomysin inhibition in 384-well plates (**Figure 3D**).

### Study of p53-HDM2 interaction and its inhibition

The data obtained with the FKBP12/FRB model system clearly shows the strong potential of Tb(III)-based, single-chain LRET biosensors for both imaging and HTS analysis of PPIs. We further evaluated the potential of these sensors by measuring the inhibition of the interaction between p53 and HDM2. As a tumor suppressor, p53 plays a crucial role in human cancer. Its activity is controlled through a negative feedback mechanism by HDM2^*37*^. The small molecule inhibitor of p53/HDM2 interaction, Nutlin-3, was identified in a screening campaign and represents one of the early successes of discovery efforts to find drugs that target PPIs^*37*^. We replaced the FRB domain in our original biosensors with the N-terminal 92 amino acids of p53 and replaced the FKBP12 domain with the N-terminal 128 residues of HDM2. Again, we prepared sensor constructs with 10, 20 and 30 nm ER/K linkers, and we stably transformed HeLa cells with the constructs for evaluation with Nutlin-3 as a positive control.

We first examined the p53/HDM2 sensor performance microscopically. Stably transformed HeLa cells were incubated with medium containing either 10 µM DMSO (negative control), or 10 µM Nutlin-3 (positive control) at 37°C for 90 min. After incubation with cell permeable TMP-Lumi4-R_9_, steady-state images of GFP fluorescence and time-gated images of Tb (III) luminescence and Tb(III)-to-GFP sensitized emission were acquired separately. A representative set of images obtained from cells expressing the p53/HDM2 sensor with a 20 nm ER/K linker clearly show a reduction in the LRET/Tb signal in cells with Nutlin-3 (**Figure 4A**). Quantitative image analysis once again showed that the maximum difference between On- and Off-states of the sensors increased with ER/K linker length; the mean decrease in LRET/Tb due to Nutlin-3 inhibition was measured to be 40%, 73%, and 84% in cells expressing sensors with 10, 20 and 30 nm linkers, respectively (**Figure 4B**). In addition, analysis of a time-lapse image sequence shows that inhibition occurs rapidly, with near maximal sensor response occurring within minutes of Nutlin-3 addition to cell culture.

**Figure 4.**
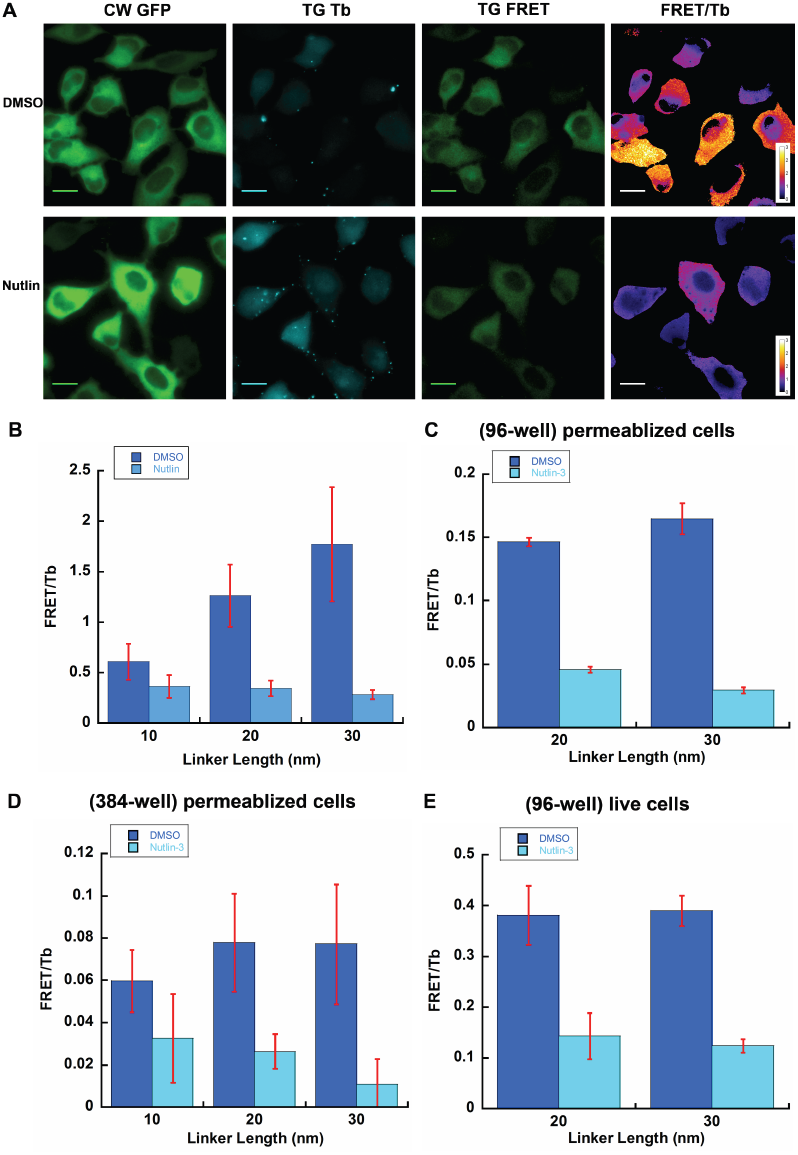
Large reductions in LRET/Tb are observed microscopically and in multiwell plates when Nutlin-3 inhibits p53/HDM2 interaction. **A.** HeLa cells stably expressing p53(1-92)-eDHFR-(ER/K)_20_-GFP-HDM2(1-128) were imaged as in **Figure 2**. Representative images show diminished LRET/Tb ratio in cells that were incubated with media containing Nutlin-3 (10 µM). **B**. Bar graphs depict the mean, pixel-wise LRET/Tb ratio measured in regions of interest drawn within cells. Values given are averaged from 10 or more cells for each condition. **C-E**. HeLa cells expressing p53/HDM2 sensor were grown in 96-well (**C, E**) or 384-well (**D**) plates. **C, D.** Time-gated measurements were obtained following overnight induction of biosensor expression with doxycycline and addition of lysis buffer containing TMP-TTHA-cs124 (50 nM) and Nutlin-3 (10 µM, positive controls) or DMSO (negative controls). **E.** Cells expressing biosensor were incubated in medium containing cell-permeable Tb complex, TMP-Lumi4-R_9_ (10 µM, RT, 30 min), washed 1X PBS and incubated in PBS containing either DMSO (negative controls), or Nutlin-3 (10 µM, positive controls). Time-gated signals were then recorded. Bar graphs depict mean LRET/Tb ratio measured for positive controls (n = 16) and for negative controls (n = 16 for **D, E** or 8 for **C**). Error bars, SEM.

We also performed the inhibition assays with permeabilized cells in both 96-well and 384-well plates using similar conditions to those used with the FKBP/FRB biosensors (**Figure 4C-D**). Following overnight induction of biosensor expression with doxycycline, lysis buffer containing TMP-TTHA-cs124(Tb) (final conc., 50 nM) was added to wells. Nutlin-3 (final conc., 10 µM) was also added with lysis buffer to positive control wells. Z’ factor values were calculated to determine data quality. In all assays, with cells expressing 10 nm linker biosensor, Z’ factor values were very low due to the low response from positive controls, in other words, trivial FRET changes occurred after incubation with Nutlin-3. However, Z’ factors ≥ 0.67, were obtained in 96-well plate assays with cell expressing 20 nm, 30 nm linker biosensor. In 384-well plate assays, Z’ factor values were negative in all cases.

### Plate reader analysis of p53-HDM2 inhibition in live cells

The ability to robustly detect PPIs or their inhibition in mammalian cell culture following cell permeabilization offers distinct benefits for drug discovery and HTS. First, no protein purification is required, and it may be possible to design expression constructs where one of the affinity partners is a transmembrane protein. Second, because the sensors are expressed directly in mammalian cells, PPIs that depend on phosphorylation or other post-translational modifications may be assessed. Finally, the assay is simple, requiring only addition of lysis buffer with detection reagent, no wash steps and immediate readout. These capabilities critically require TGL detection of lanthanide-based FRET as well as a single-chain biosensor design. For example, only ∼8,000 cells are present in a single well of a 384-well plate with a solution volume of 50 µL. If we assume a cell volume of 3 pL and a moderate biosensor expression level such that effective cellular concentration is 5 µM, then only sub-picomolar amounts of protein are present in a well and the sensor concentration following cell lysis is in the low-nanomolar range. These concentrations are below the detection limits of conventional FRET and far below the *K*_*D*_ of most relevant PPIs. Consequently, the affinity elements must be tethered to one another and the high sensitivity of TGL detection is needed.

While PPI detection following cell permeabilization offers substantial benefits, the ability to detect PPI changes within intact, live cells could offer more biologically relevant insights as it would allow for PPI analysis in the presence of other cellular factors. Moreover, HTS assays within live cells would further assess the ability of drugs to cross the plasma membrane and their inhibition or activation characteristics within the cellular milieu. We evaluated the performance of our sensors in live cells in 96-well plates using the same, cell permeable TMP-Lumi4-R_9_ complex that we used for microscopic imaging. After overnight induction in medium containing doxycycline, HeLa cells stably expressing single-chain p53/HDM2 affinity biosensors with 20 nm or 30 nm ER/K linker were washed and incubated in medium containing 10 µM TMP-Lumi4-R_9_ at room temperature for 30 min. Following DPBS wash, PBS buffer solution containing either 10 µM DMSO (negative control), or 10 µM Nutlin-3 (positive control) was added to wells and left at room temperature for 40 min. Time-gated Tb and Tb-sensitized FRET signals were then recorded. We calculated a Z’ factor of 0.50 for cells expressing 30 nm linker biosensor (**Figure 4E**). This good Z’ factor with manual plate preparation advocates the application of the 30 nm linker single-chain biosensor in HTS assay for live cell PPIs interaction or inhibition hit selection.

## CONCLUSIONS

Single-chain LRET biosensors have a number of unique benefits for live-cell imaging and cell-based screening of PPIs. Extraordinary dynamic range stems from time-gated detection of Tb(III)-to-GFP LRET that eliminates non-specific fluorescent background and from incorporation of an alpha-helical, ER/K linker that maintains Tb(III) donors and GFP acceptors far apart when the sensor is in the open configuration. These features enable dynamic visualization of PPIs in cells with TGL microscopy, robust detection of PPIs or their inhibition within intact cells grown in 96-well plates, or high throughput detection in cell lysates in 384-well plates. In principle, it should be possible to detect interactions between a membrane protein and a cytosolic protein or between proteins that are otherwise difficult or impossible to purify. Moreover, Tb(III) can sensitize differently colored acceptors, offering the potential for multiplexed imaging or analysis. Taken together, the results presented here show that Tb(III)-based LRET biosensors offer a versatile platform technology for interrogating PPIs and their function in live cells.

## EXPERIMENTAL METHODS

### Materials

Dulbecco’s modified eagle medium with 1g/L glucose (DMEM, 10-014CV), Dulbecco’s modified eagle medium with 4.5g/L glucose (DMEM, 10-013CV), Dulbecco’s phosphate buffer saline (DPBS, 21-030 and 21-031), 0.25% trypsin/2.21 mM EDTA and 0.05% trypsin/2.21 mM EDTA (25-053-Cl) were purchased from Corning cellgro^®^. MEM non-essential amino acid (11140), DMEM (without phenol red, 21063), HEPES (15630-080) and Lipofectamine 2000 (11668-027) were purchased from Invitrogen™. FBS (S11150) was purchased from Atlanta Biologicals. Hygromycin (sc-29067) and Nutlin-3 (sc45061) were purchased from Santa Cruz Biotechnology. BSA (700-107P) was purchased from Gemini Bio-products. Rapamycin (553211-500UG) was purchased from Millipore. Ascomycin (11309) was purchased from Cayman Chemical. NADPH (N0411) and doxycycline (D9891) were purchased from Sigma. DMSO (D128-500) was purchased from Fisher Chemical. Patent V blue sodium salt (21605) was purchased from Fluka. Clontech In-Fusion^®^ Cloning Kit (638909) was purchased from Takara. All enzymes and buffers used in cloning were purchased from New England Biolabs.

### Luminescent Tb(III) complexes

Heterodimers of trimethoprim linked to luminescent Tb(III) complexes (TMP-cs124-TTHA,^*19, 34*^ and TMP-Lumi4,^*19*^) and a cell permeable variant conjugated to oligoarginine (TMP-Lumi4-R_9_^*32*^) were prepared as previously reported.

### Plasmids

All DNA constructs were sequenced by the UIC Research Resources Center (RRC).

### pPBH-TRE_tight_-FRB-eDHFR-(ER/K)_10nm_-EGFP-FKBP12

was prepared via subcloning from a pUC67 plasmid that contained the ORF, FRB-eDHFR-(ER/K)_10nm_-EGFP-FKBP12. GenScript, Inc. prepared the source vector using plasmid DNA that contained the fragments EGFP-FKBP12 and FRB-eDHFR (see Yapici^*38*^) and synthesized DNA encoding an ER/K linker of length 10 nm with the sequence of 5’ – GAA GAG GAA GAG AAA AAA AAA CAG CAG GAA GAG GAA GCA GAA AGG CTG AGG CGT ATT CAA GAA GAA ATG GAA AAG GAA AGA AAA AGA CGT GAA GAA GAC GAA AAA CGT CGA AGA AAG GAA GAG GAG GAA AGG CGG ATG AAA CTT GAG ATG GAA GCA AAG AGA AAA CAA GAA GAA GAA GAG AGA AAG AAA AGG GAA GAT GAT GAA AAA CGC AAG AAG AAG. The ORF was inserted into the pPBH-TRE_tight_ vector between KpnI site and NheI site to give pPBH-TRE_tight_-FRB-eDHFR-(ER/K)_10nm_ -EGFP-FKBP12.

### pPBH-TRE_tight_-FRB-eDHFR-(ER/K)_30nm_-EGFP-FKBP12

A 630 bp (ER/K)_30nm_ linker fragment (sequence reported in Sivaramakdrishnan and Spudich^*30*^) was synthesized and cloned into pUC57 vector by GenScript, Inc. The genes encoding FRB-eDHFR, (ER/K)_30nm_, EGFP-FKBP12 were subcloned from plasmids pPBH-TREtight-FRB-eDHFR-(ER/K)_10nm_-EGFP-FKBP12 and (ER/K)_30nm_ in pUC57 to pPBH-TREtight to generate pPBH-TREtight-FRB-eDHFR-(ER/K)_30nm_-EGFP-FKBP12. A 753 bp fragment encoding FRB-eDHFR was prepared by PCR from pPBH-TREtight-FRB-eDHFR-(ER/K)_10nm_-EGFP-FKBP12 using the primers 5’-ACT CTG CAG TCG ACG GTA CCA TGA TCC TCT GGC ATG AGA TGT GGC -3’ (coding strand) and 5’-TCG GAT CCT CCG CTT CCC CGC CG -3’ (non-coding strand). A 630 bp fragment encoding (ER/K)_30nm_ was prepared by PCR from (ER/K)_30nm_ in pUC57 using the primers 5’-AAG CGG AGG ATC CGA AGA GGA GGA GAA AAA GAA GGA -3’ (coding strand) and 5’-CCA GAG CCA CCG GTT CTC TGT TTT CGC TCT GC -3’ (non-coding strand). A 1041 bp fragment encoding EGFP-FKBP12 was prepared by PCR from pPBH-TREtight-FRB-eDHFR-(ER/K)_10nm_-EGFP-FKBP12 using the primers 5’-AAC CGG TGG CTC TGG CAT GGT GAG CA -3’ (coding strand) and 5’-ATG CGG CCG CGC TAG-3’ (non-coding strand). These 3 fragments were inserted between the KpnI site and the NheI site in pPBH-TREtight by Clontech In-Fusion^®^ Cloning Kit to get pPBH-TREtight-FRB-eDHFR-(ER/K)_30nm_-EGFP-FKBP12.

### pPBH-TRE_tight_-FRB-eDHFR-(ER/K)_20nm_-EGFP-FKBP12

The (ER/K)_20nm_ linker is comprised of the first 396 bp of the (ER/K)_30nm_ linker. The genes encoding FRB-eDHFR, (ER/K)_20nm_, EGFP-FKBP12 were subcloned from plasmids pPBH-TREtight-FRB-eDHFR-(ER/K)_10nm_-EGFP-FKBP12 and (ER/K)_30nm_ in pUC57 to pPBH-TREtight to generate pPBH-TREtight-FRB-eDHFR-(ER/K)_20nm_-EGFP-FKBP12. A 753 bp fragment encoding FRB-eDHFR was prepared by PCR from pPBH-TREtight-FRB-eDHFR-(ER/K)_10nm_-EGFP-FKBP12 using the primers 5’-ACT CTG CAG TCG ACG GTA CCA TGA TCC TCT GGC ATG AGA TGT GGC -3’ (coding strand) and 5’-TCG GAT CCT CCG CTT CCC CGC CG -3’ (non-coding strand). A 396 bp fragment encoding (ER/K)_20nm_ was prepared by PCR from (ER/K)_30nm_ in pUC57 using the primers 5’-AAG CGG AGG ATC CGA AGA GGA GGA GAA AAA GAA GGA -3’ (coding strand) and 5’-CCA GAG CCA CCG GTC TCT TCC TTG GCC TTT TTC TCC TGC -3’ (non-coding strand). A 1041 bp fragment encoding EGFP-FKBP12 was prepared by PCR from pPBH-TREtight-FRB-eDHFR-(ER/K)_10nm_ -EGFP-FKBP12 using the primers 5’-GAA GAG ACC GGT GGC TCT GGC ATG GTG AGC A -3’ (coding strand) and 5’-ATG CGG CCG CGC TAG-3’ (non-coding strand). These 3 fragments were inserted between the KpnI site and the NheI site in pPBH-TRE_tight_ by Clontech In-Fusion^®^ Cloning Kit to get pPBH-TRE_tight_-FRB-eDHFR-(ER/K)_20nm_ -EGFP-FKBP12.

### pPBH-TRE_tight_-p53(1-92)-eDHFR-(ER/K)_n_-EGFP-HDM2(1-128)

The genes encoding p53(1-92), FRB-eDHFR-(ER/K)_n_-EGFP (n=10 nm, 20nm or 30 nm), and HDM2 (1-128) were subcloned from plasmids p53-GFP, pPBH-TRE_tight_-FRB-eDHFR-(ER/K)_n_-EGFP-FKBP12 (n = 10 nm, 20nm or 30 nm) and pCMV-HDM2(C464A) to pPBH-TRE_tight_ to generate pPBH-TRE_tight_-p53(1-92)-eDHFR-(ER/K)_n_-EGFP-HDM2(1-128). A 276 bp fragment encoding p53 (residues 1-92) was prepared by PCR from p53-GFP using the primers 5’-ACT CTG CAG TCG ACG GTA CCA TGG AGG AGC CGC AGT CA -3’ (coding strand) and 5’-CCA GAT CCG GGC CAG GAG GGG G -3’ (non-coding strand). Fragments of length 1446 bp, 1620 bp, or 1845 bp that encoded eDHFR-(ER/K)_n_-EGFP where n equaled 10 nm, 20nm or 30 nm, respectively, were prepared by PCR from pPBH-TREtight-FRB-eDHFR-(ER/K)n-EGFP-FKBP12 (n=10 nm, 20nm or 30 nm) using the primers 5’-CTG GCC CGG ATC TGG AGG ATC TGG AAT CAG TC -3’ (coding strand) and 5’-TTG CAC ATT CGA GAT CTG AGT CCG GAC TTG TA -3’ (non-coding strand). A 384 bp fragment encoding HDM2 (residues 1-128) was prepared by PCR from pCMV-HDM2(C464A) using the primers 5’-ATC TCG AAT GTG CAA TAC CAA CAT GTC TGT ACC -3’ (coding strand) and 5’-ATG CGG CCG CGC TAG CCT ATT CAA GGT GAC ACC TGT TCT CAC TC -3’ (non-coding strand). These 3 fragments were inserted between the KpnI site and the NheI site in pPBH-TRE_tight_ using Clontech In-Fusion^®^ Cloning Kit to obtain pPBH-TRE_tight_-p53(1-92)-eDHFR-(ER/K)_n_-EGFP-HDM2(1-128) (n=10 nm, 20nm or 30 nm).

### Stable expression of biosensor plasmids

All FRB/FKBP12 biosensor plasmids were transfected to NIH 3T3 cells, while all p53/HDM2 biosensors were transfected to HeLa cells. Cells were grown to 70-80% confluency in a sterile 10 cm dish. The cells were transfected with 12 μg of biosensor plasmid DNA and the recombination helper plasmid pSPB-Transposase with a Lipofectamine:plasmid ratio of 2.5µL:1µg per plasmid. Plasmid and Lipofectamine solutions were first prepared in separate microcentrifuge tubes in OptiMEM I with a total volume of 1.5 mL. After 5 minutes of incubation at room temperature, the solutions were mixed and kept at room temperature for an additional 20 minutes. The media in 10 cm dish was aspirated and the Lipofectamine + plasmids solution was added into it. The cells were incubated with the solution for 4 hours at 37 °C with 5% CO_2_, and then the solution was replaced with 10 mL of fresh DMEM(+) (DMEM supplied with 15 mM HEPES, 10% FBS and 100 mg/mL Hygromycin). The transfections were confirmed with microscopy and/or flow cytometry by using the GFP emissions.

### Probe delivery for TGLM

Cells were trypsinized and seeded at 20,000 cells/well in an 8-well chambered coverglass (Nunc™, 12-565-470) with fresh DMEM (+) containing 100 ng/mL doxycycline to induce the expression of proteins and incubated at 37 °C and 5% CO_2_ overnight. For FRB/FKBP12 stable transfected cell lines, on the following day the cells were washed twice with DPBS (+Ca/+Mg), 100 μL of TMP-Lumi4-R9 (12 μM in DMEM without phenol red) was added, and the cells were incubated for 15 min at room temperature. Cells were washed again with DPBS (+Ca/+Mg) and 150 μL of DMEM without phenol with Rapamycin (1 μM, 1% DMSO) was added. Control wells received DMEM without phenol red with DMSO (1%) but without rapamycin. The cells were incubated for 15 min at 37 °C and 5% CO_2_. Immediately prior to microscope imaging, 20μL of 10 mM patent blue V solution (final concentration: 1 mM) was added to quench extracellular luminescence from non-specifically adsorbed probe. To obtain the time-lapse images of FRB/FKBP12 interaction, cells were first loaded with TMP-Lumi4-R_9_, washed, immersed in DMEM (without phenol red) with patent blue (1 mM) and then rapamycin was added (final concentration: 2 µM).

HeLa cells stably expressing the p53/HDM2 biosensor were seeded into chambered coverglass (20,000 cells/well) and incubated overnight in DMEM with 100 ng/mL to induce protein expression. On the day after seeding, the cells were incubated with DMEM without FBS containing Nutlin-3 (final conc. 10 µM) or vehicle (DMSO). for 90 min at 37 °C and 5% CO_2_. The cells were washed twice with DPBS (+Ca/+Mg), 100 μL of TMP-Lumi4-R9 (12 μM in DMEM without phenol red) was added, and the cells were incubated for 20 min at room temperature. Cells were washed again with DPBS (+Ca/+Mg) and 150 μL of patent blue V solution (1 mM in DMEM without phenol red, containing 10 µM Nutlin-3) was added to the sample well for microscope imaging.

### Time-gated Luminescence Microscopy and image processing

Time-gated luminescence images were acquired using a previously described epi-fluorescence microscope (Axiovert 200, Carl Zeiss, Inc.).^*39, 40*^ For each time-gated image acquisition, the signal from multiple excitation/emission events was accumulated on the ICCD sensor and read out at the end of the camera frame. The UV LED pulse width and pulse period, the intensifier delay time and on-time, the camera frame length (66.67 ms – 2 s) and the intensifier gain voltage could be varied independently. The source/camera timing parameters were the same for all of the time-resolved images and data presented here: excitation pulse width, 1500 μs, pulse period, 3000 μs, delay time, 10 μs, intensifier on-time, 1480 μs. All data reported here was acquired at a gain of 833 V. The camera control software enabled summation of multiple frames to yield a single composite .TIFF image with a bit depth equal to 1024 multiplied by the number of frames. All images reported here were summations of four frames (bit depth, 4096), and a feature of the camera control software was enabled that removes large variations in signal resulting from ion-feedback noise of the intensifier. The emission filters and all detector timing and gain parameters for each figure in the text are provided in Table S1.

Raw, 12-bit images were imported into NIH ImageJ (v1.42q) for all processing operations including cropping, contrast adjustment, and quantitative analysis.^*41*^ For each channel, 20 dark frames and 20 bright field images were stacked, converted to 32 bits, and median-filtered (radius 1), and each stack was averaged. The flat-field average was divided by the mean intensity of its central nine pixels to generate a normalized flat-field image. For each sample image, a median filter (radius 1) was applied and the master dark frame was subtracted. The resulting image was then divided by the normalized, master flat-field image, and the mean value of the detector offset was added back to the image. For ratiometric images and measurements, a binary mask was created by first averaging a series of GFP images and then applying a threshold to highlight only regions exhibiting signal. The mask was applied to background-subtracted time-gated FRET images, and the FRET images were then divided by the GFP or Tb image. Intensity-modulated ratiometric displays were generated using the Fire lookup table in ImageJ and a color lookup table was applied.

### Multi-well plate assays

#### Rapamycin stimulation assay with permeabilized mammalian cells

NIH3T3 fibroblasts stably expressing pPBH-TRE_tight_-FRB-eDHFR-(ER/K)_n_-EGFP-FKBP12 (n=10nm, 20nm or 30 nm) were seeded into multiwell plates at a density of 1.6 × 10^5^ cells/mL (250 µL for 96-well plate, 50 µL for 384-well plate) and incubated (37 °C, 5% CO_2_) for 24 h in culture medium containing 100 ng/mL doxycycline. For the titration assay, growth media was removed carefully with a hand pipette, and 50 µL lysis buffer (5 μM NADPH, 0.1% BSA, 0.1% Triton X-100 in DPBS) containing TMP-Lumi4 (50 nM) and rapamycin (0.47 – 5 μM) was added into the wells. For single-point assays, growth media in the wells were discarded carefully and lysis buffer with TMP-cs124-TTHA (25 nM) and rapamycin (1 µM rapamycin) was added into the wells (50 µL for 96-well plates, 30 µL for 384-well plates). The plates were kept at room temperature in dark for 15 min prior to the first measurement. Negative control wells contained lysis buffer with Tb(III) complex but no cells.

#### Ascomycin inhibition assay with permeabilized mammalian cells

NIH 3T3 fibroblasts stably expressing pPBH-TRE_tight_-FRB-eDHFR-(ER/K)_n_-EGFP-FKBP12 (n=10nm, 20nm or 30 nm) were prepared as above. Lysis buffer containing TMP-cs124-TTHA (50 nM) and ascomycin (final conc. 0.02 µM – 40 µM for titration assay; 20 µM for single point inhibition assay) was added into wells (50 µL for 96-well plate, 30 µL for 384-well plate). The plate was kept at room temperature in dark for 20 minutes prior to the first measurement. Subsequent measurements were recorded every 10 minutes up to 2.5 hours. Negative control wells contained cells without protein expression, but the same lysis buffer as sample wells.

#### Nutlin-3 inhibition assay with permeabilized mammalian cells

pPBH-TRE_tight_-p53(1-92a.a.)-eDHFR-(ER/K)_n_-EGFP-HDM2(1-128a.a.) (n=10 nm, 20 nm or 30 nm) stable transfected cells were seeded at a density of 1.6 × 10^5^ cells/well in a multi-well plate and incubated (37 °C, 5% CO_2_) for 24 h in culture medium (250 µL for 96-well plate, 50 µL for 384-well plate) containing 100 ng/mL doxycycline. The following day, for the titration assay, growth media in the wells were discarded carefully and lysis buffer (50 nM TMP-cs124-TTHA-Tb^3+^, 5 μM NADPH, 0.1% BSA, 0.1% Triton X-100, and Nutlin-3 with 2-fold serial dilution in the range of 200 μM – 0.098 µM in DPBS solution) was added into the wells; for the inhibition assay, growth media in the wells were discarded carefully and lysis buffer (50 µL for 96-well plate, 30 µL for 384-well plate) containing 50 nM TMP-cs124-TTHA-Tb^3+^, 5 μM NADPH, 0.1% BSA, 0.1% Triton X-100, 10 µM Nutlin-3 in DPBS solution was added into the wells. Then the plate was kept at room temperature in dark for 20 minutes and the first measurement was taken afterwards. Negative control wells contained cells without protein expression, but the same lysis buffer as sample wells.

#### Nutlin-3 inhibition assay with live mammalian cells

HeLa cells stably expressing pPBH-TRE_tight_-p53(1-92)-eDHFR-(ER/K)_n_-EGFP-HDM2(1-128) (n=20 nm or 30 nm) were grown in 96-well plates and protein expression was induced in the same manner described above for NIH 3T3 cells. The cells were incubated at room temperature for 30 min in DMEM without phenol red containing TMP-Lumi4-R_9_ (10 µM). The media were removed, cells were washed twice with DPBS and DMEM (without phenol red) containing Nutlin-3 (10 µM, 1% DMSO) or vehicle was added into the wells. The plate was kept at room temperature in dark for 40 minutes prior to the first measurement. Negative control wells contained cells without protein expression, but the same solutions as sample wells.

## ACKNOWLEDGMENTS

We thank Lumiphore, Inc. for providing Lumi4-Tb^®^ and TMP-Lumi4-R_9_. Funding was provided by NIGMS (R01 GM081030).

