## Supplemental Information for "Single-chain lanthanide luminescence biosensors for cell-based imaging and screening of protein-protein interactions"

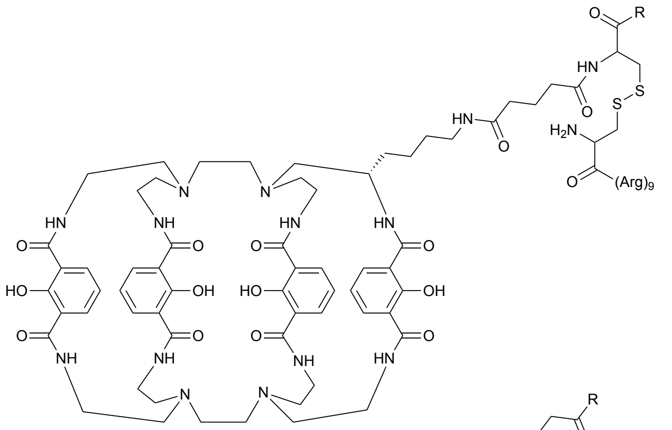

1

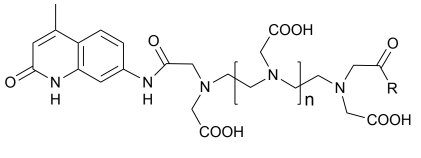

2 : n = 2

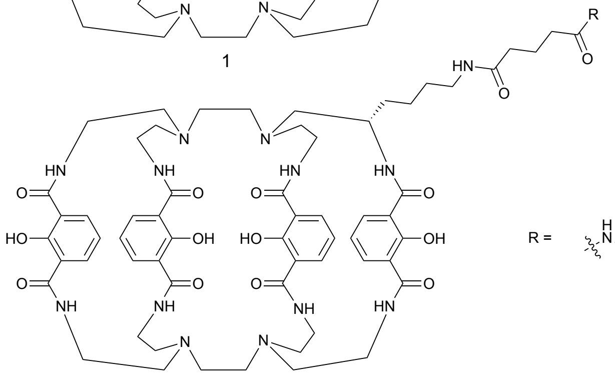

3

R = 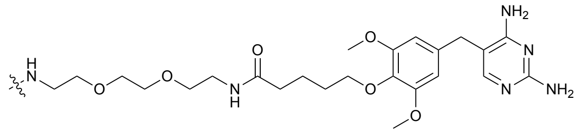

**Figure S1**. Chemical structures of TMP-Lumi4-R9 (**1**), TMP-TTHA-cs124 (**2**) and TMP-Lumi4 (**3**). Probe (**1**) was used in microscopic imaging while (**2**) and (**3**) were used in plate-reader assay.

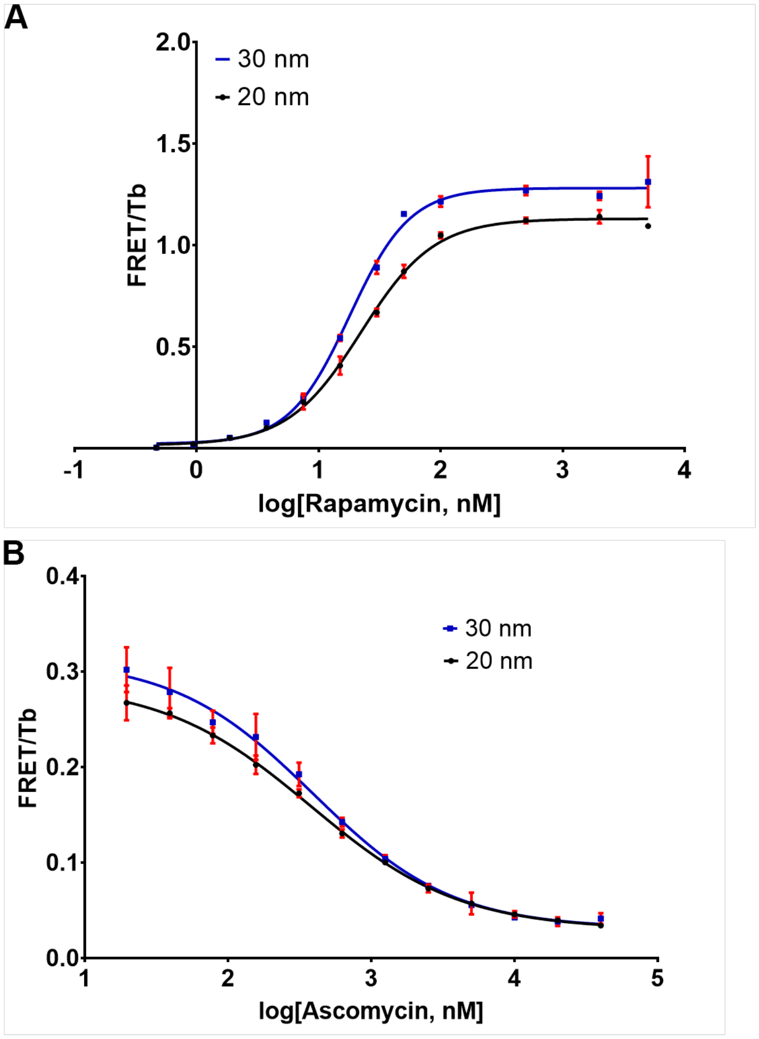


**Figure S2**. **Rapamycin and ascomycin titration assay**. NIH 3T3 cells expressing FKBP/FRB biosensors (containing 20 nm or 30 nm ER/K linkers) were seeded into 96-well plates. Following overnight incubation, cells were treated with lysis buffer containing TMP-Lumi4-Tb (50 nM) and (A) serial dilutions of rapamycin (final conc., 5 μM to 0.5 nM) or (B) 0.333 μM rapamycin and a serial dilutions of ascomycin (final conc., 40 μM to 0.02 μM).
